# Dnmt1 determines bone length by regulating energy metabolism of growth plate chondrocytes

**DOI:** 10.1101/2024.07.17.604010

**Authors:** Yuta Yanagihara, Masatomo Takahashi, Yoshihiro Izumi, Tomofumi Kinoshita, Masaki Takao, Takeshi Bamba, Yuuki Imai

**Author notes:** All correspondence to Yuuki Imai, M.D., Ph.D. Division of Integrative Pathophysiology, Proteo-Science Center, Ehime University Shitsukawa, Toon, Ehime 791-0295 JAPAN.

## Abstract

Chondrocytes differentiated from mesenchymal stem cells play a role in determining skeletal patterns by ossification. However, the mechanism by which maintenance DNA methylation in chondrocytes regulates differentiation and skeletal formation is unclear. In the Musculoskeletal Knowledge Portal, Dnmt1 was significantly associated with “Height”. Long bones in the limbs of Dnmt1-deficient (*Dnmt1^ΔPrx1^*) mice are significantly shortened due to decreased chondrocyte proliferation and accelerated differentiation. Integrated analysis of RNA-Seq and MBD-Seq revealed that in *Dnmt1^ΔPrx1^* chondrocytes reduced DNA methylation resulted in increased expression of genes related to energy metabolism and to ossification. Metabolomic analyses confirmed that levels of nearly all energy metabolites were increased in *Dnmt1^ΔPrx1^* chondrocytes. These results indicate that Dnmt1-mediated maintenance DNA methylation governs chondrocyte differentiation by regulating energy metabolism through both gene expression and modulation of metabolite supplies. Taken together, this study suggests that appropriate DNA methylation status in chondrocytes can orchestrate growth plate mineralization and subsequently determine bone length.

## Introduction

Chondrocytes are involved in formation of cartilage and play a pivotal role in the development of the skeletal system. The formation of skeletal elements, including long bones, occurs through endochondral ossification when mesenchymal stem cells differentiate into chondrocytes that then form an anlage of cartilage. The cartilage anlage serves as a scaffold for bone formation. Chondrocytes within the cartilage matrix secrete extracellular matrix components, including collagen and proteoglycans. As chondrocytes mature and undergo hypertrophy, they begin to mineralize the surrounding matrix. Ultimately, the hypertrophic cartilage is replaced by bone tissue, which contributes to longitudinal bone growth. This chondrocyte differentiation process is essential for the development, growth, and maintenance of the skeletal system ^1^.

Epigenetics involves regulation of gene expression without altering the genomic DNA sequence. During cell differentiation, epigenetic modifications such as DNA methylation, histone modifications, and chromatin remodeling dynamically regulate the accessibility of transcriptional machinery to gene loci ^2^. Epigenetic mechanisms play a crucial role in cell differentiation and development. In particular, DNA methylation maintenance is an epigenome-supporting system that reconstructs histone modifications and chromatin structure in post-mitotic cells through inheritance of DNA methylation ^3^. The DNA methylation maintenance system comprises two key molecules: Ubiquitin-like containing PHD and RING finger domains 1 (Uhrf1) and DNA methyl transferase 1 (Dnmt1). During DNA replication, Uhrf1 recognizes hemimethylated DNA and recruits Dnmt1, which methylates the replicated DNA strand. DNA methylation in the vicinity of the transcription start site is known to suppress gene expression ^4^.

As we previously reported, Uhrf1 regulates limb growth through control of gene expression ^5^. Uhrf1 has various domains and diverse functions, including maintaining DNA methylation as well as de novo methylation and genomic DNA stability ^6, 7^. However, the relationship between bone growth and mechanisms of Dnmt1, the main DNA methyltransferase involved in DNA methylation maintenance, remained unclear. Dnmt1 was previously reported to be involved in knee osteoarthritis (OA) and mineralization of periodontal ligament cells and mesenchymal cells ^8, 9, 10, 11^, but the function of Dnmt1 in skeletal formation, particularly in chondrocytes, is not known. Here we show that Dnmt1 regulates not only gene expression but also energy metabolism that is involved in mineralization of cartilage and consequent determination of bone length.

## Results

### Lack of growth of long bones in *Dnmt1^ΔPrx1^* mice after birth

To elucidate the role of Dnmt1 in the skeletal system, we first used the Musculoskeletal Knowledge Portal (MSK-KP) to assess the gene-level association of single nucleotide polymorphisms in the DNMT1 gene locus. The most significant correlation was with ‘Height’ (P value < 4.2e^-35^; Figure 1A). Therefore, we generated mice with conditional knockout of *Dnmt1* that was specific to limb mesenchymal cells (*Dnmt1^ΔPrx1^*) by crossing *Prrx1*-Cre mice with *Dnmt1 floxed* mice and analyzing the effects of Dnmt1 knockout on skeletal formation. Skeletal preparations from neonatal control (*Dnmt1^flox^*) mice and *Dnmt1^ΔPrx1^* mice showed that both genotypes had mineralized bone and cartilage tissue in the limbs (Figure 1B). However, the joints of *Dnmt1^ΔPrx1^* mice had a poorly formed cartilage matrix at 1 week after birth (Figure 1C). The difference in bone length became more pronounced as the mice aged, with *Dnmt1^ΔPrx1^*mice having bone length that was approximately 60% (Figure 1D, S1A, B), 53% (Figure S1C), and 43% (Figure 1E, F, S1A, D, E) that of *Dnmt1^flox^* mice at 1, 2, 3, and 6 weeks of age, respectively. These data indicated that Dnmt1 plays more important roles in longitudinal bone growth in the postnatal rather than embryonic developmental stage.

**Figure 1.**
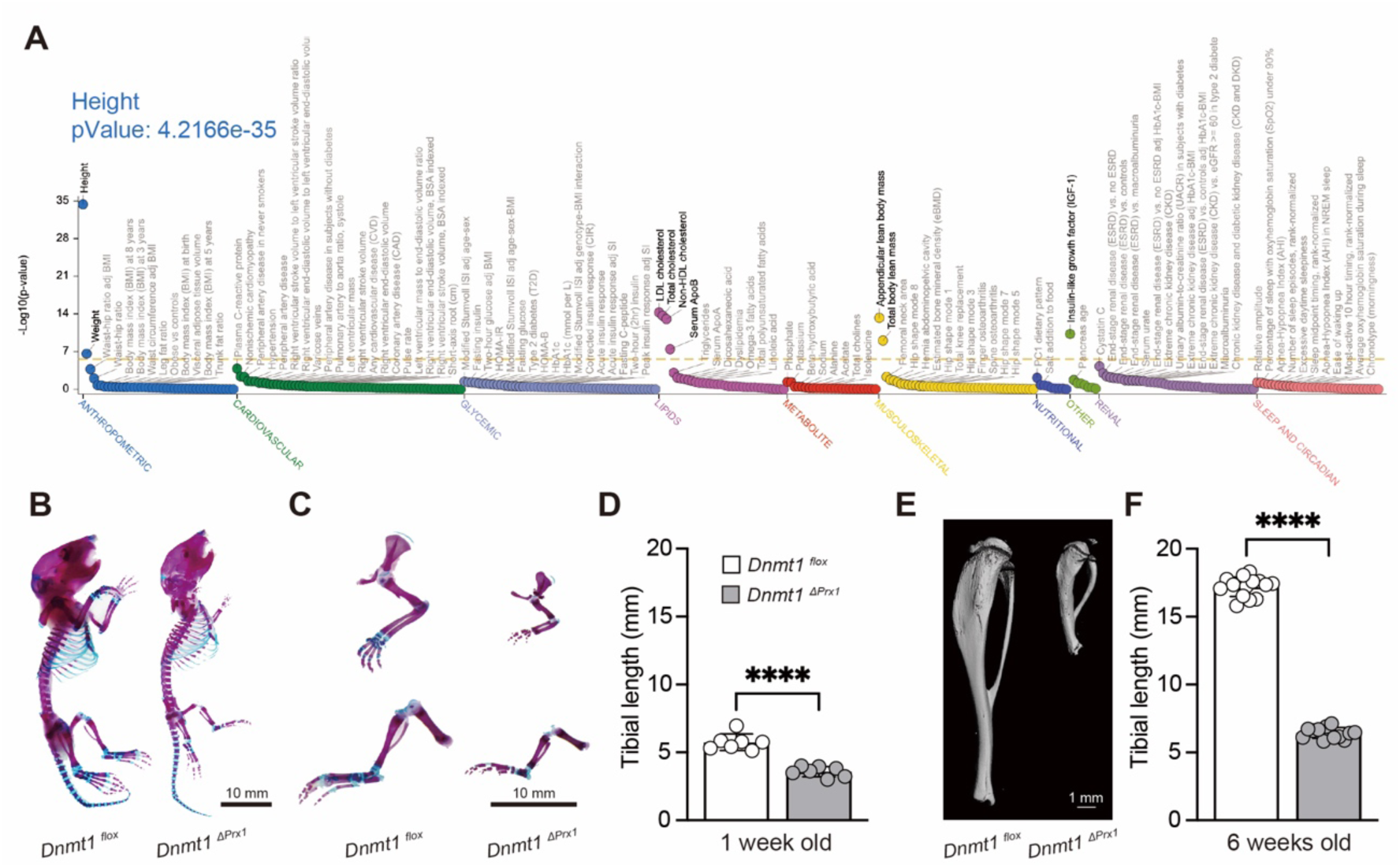
Mice with Dnmt1 knockout specific to limb mesenchymal cells exhibit significant limb shortening. (A) Gene-level association of single nucleotide polymorphisms in the DNMT1 gene locus obtained from the Musculoskeletal knowledge portal (MSK-KP). (B) Skeletal preparations (Alcian Blue/Alizarin Red) of neonatal *Dnmt1^flox^* and *Dnmt1^ΔPrx1^* mice. (C) Higher magnification images of skeletal preparations of forelimbs (upper panels) and hindlimbs (lower panels). (D) Length of tibiae from 1-week-old *Dnmt1^flox^* (n=7) and *Dnmt1^ΔPrx1^*(n=7) mice. (E) µCT images of tibiae from 6-week-old *Dnmt1^flox^* and *Dnmt1^ΔPrx1^* mice. (F) Length of tibiae from 6-week-old *Dnmt1^flox^* (n=15) and *Dnmt1^ΔPrx1^*(n=13) mice. (B, C) Representative data from at least three individual mice are shown. All data are mean ± s.d. **** *p* < 0.0001 (unpaired *t* test).

### Suppressed proliferation and accelerated differentiation in *Dnmt1^ΔPrx1^*growth plates

Growth plate cartilage is important for longitudinal bone elongation. Here we performed histological analyses to determine whether Dnmt1 is expressed in growth plate cartilage. Immunohistochemical staining revealed that expression of both Dnmt1 and Uhrf1 localized to chondrocytes in the proliferative zone of control *Dnmt1^flox^* mice. In contrast, *Dnmt1^ΔPrx1^*mice had significantly lower Uhrf1 expression relative to *Dnmt1^flox^*mice, and essentially no Dnmt1 expression (Figure 2A, B). These results indicate that Dnmt1 functions mainly in the proliferative chondrocytes of growth plate cartilage and that Prrx1-Cre-mediated conditional knockout effectively reduced Dnmt1 expression in the limbs. Alcian Blue staining of cartilage tissue revealed that the proliferative cartilage area in the proximal tibia was significantly smaller in *Dnmt1^ΔPrx1^* mice compared to *Dnmt1^flox^* mice at 1 week of age (Figure 2C, D). The population of bromodeoxyuridine (BrdU)-positive cells in growth plate cartilage was significantly lower in *Dnmt1^ΔPrx1^*mice than control *Dnmt1^flox^*mice (Figure S2A, B) while the hypertrophic cartilage area and mineralized area were significantly wider (Figure 2C, D, E). At 6 weeks of age, *Dnmt1^ΔPrx1^* mice exhibited a loss of growth plates and trabecular bones, and a marked delay in the formation of secondary ossification centers in the proximal tibia (Figure 2F, G). The control *Dnmt1^flox^* mice had slow progression of growth plate calcification, and sequential μCT images did not exhibit trabecular bone until the mice were 6 weeks old (Figure S2C, upper panel). On the other hand, *Dnmt1^ΔPrx1^* mice had significant calcification of both growth plates and trabecular bones beginning as early as 1 week of age, and calcified trabecular structures had disappeared when the mice were 6 weeks old (Figure S2C, lower panel). Trabecular bone formation is dependent on proper osteoclast/chondroclast and osteoblast function. As such, we carried out immunohistochemical staining for osteoclast and osteoblast markers using tibiae from 2-week-old mice. The area of Osterix (Sp7)-positive staining did not differ between *Dnmt1^flox^* mice and *Dnmt1^ΔPrx1^* mice, but the area of Cathepsin K (Ctsk)-positive osteoclasts was significantly larger in *Dnmt1^ΔPrx1^* mice compared to *Dnmt1^flox^*mice (Figure S2D, E). These results revealed that *Dnmt1^ΔPrx1^* mice have reduced proliferative activity and increased calcification, even in a proliferative zone, that was followed by abnormally accelerated calcified bone/cartilage resorption resulting in the disappearance of growth plates when the mice were 6 weeks old.

**Figure 2.**
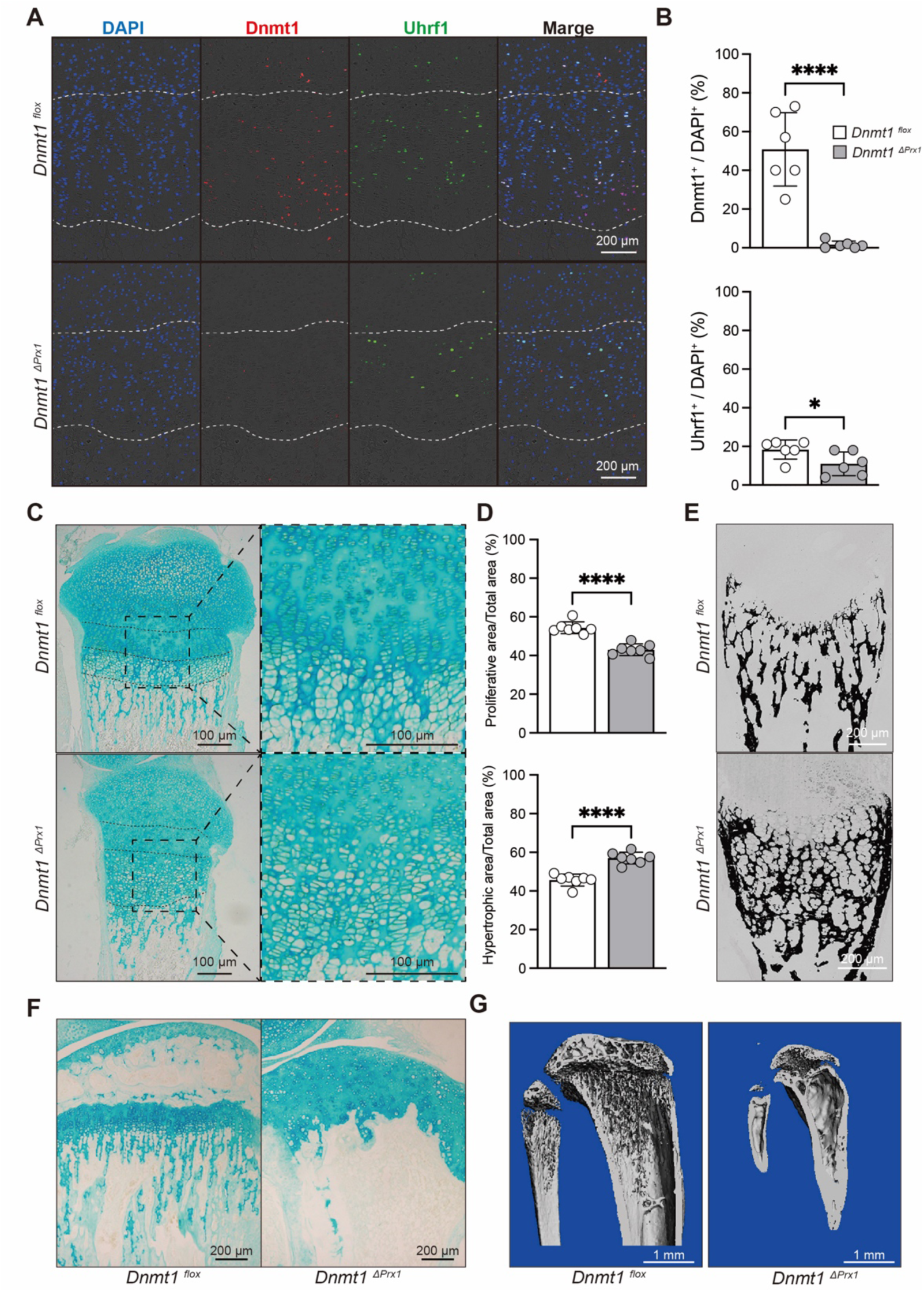
Narrowed proliferative cartilage layer and promotion of chondrocyte hypertrophy and mineralization associated with Dnmt1 deficiency. (A) Immunohistochemistry for Dnmt1 (Red), Uhrf1 (Green) and DAPI (Blue) in proximal tibiae from 1-week-old mice. (B) Quantification of Dnmt1 and Uhrf1 expression in growth plate chondrocytes from *Dnmt1^flox^* (n=6) and *Dnmt1^ΔPrx1^*(n=6) mice. (C) Alcian Blue staining of proximal tibiae from 1-week-old mice; right panels show a higher magnification of the boxed areas in the left panels. (D) Quantification of the area of proliferative and hypertrophic cartilage layers in proximal tibiae from *Dnmt1^flox^*(n=7) and *Dnmt1^ΔPrx1^* (n=7) mice. (E) Von Kossa staining of proximal tibiae from 1-week-old mice. (F) Alcian Blue staining of proximal tibiae from 6-week-old mice. (G) µCT images of tibiae from 6-week-old mice. (E-G) Representative data from at least three individual mice are shown. All data are mean ± s.d. **** *p* < 0.0001 (unpaired *t* test).

### Dnmt1 deficiency accelerates chondrocyte mineralization

To investigate the molecular basis for the *Dnmt1^ΔPrx1^*phenotype, we evaluated the differentiation of primary cultured chondrocytes obtained from neonatal *Dnmt1^flox^* and *Dnmt1^ΔPrx1^*mice. Gene expression levels of *Dnmt1* in *Dnmt1^ΔPrx1^* chondrocytes were significantly lower regardless of the presence or absence of Bone morphogenetic protein 2 (BMP2) (Figures S3A). Levels of Dnmt1 protein were also significantly lower (Figures S3B, C) in *Dnmt1^ΔPrx1^* chondrocytes relative to *Dnmt1^flox^* chondrocytes. Along with the Dnmt1 deficiency, gene expression levels of *Uhrf1* and *Ten-eleven translocation* (*Tet1/2/3*) DNA demethylases were also decreased in *Dnmt1^ΔPrx1^* mice (Figure S3A). Proliferation of chondrocytes derived from *Dnmt1^ΔPrx1^* mice was significantly slower than that for *Dnmt1^flox^* chondrocytes (Figure S3D, E). Moreover, Alcian Blue staining indicated a notable reduction in cartilage matrix synthesis in *Dnmt1^ΔPrx1^* chondrocytes (Figures 3A, B). On the other hand, consistent with *in vivo* observations (Figures 2E), mineralization was accelerated in *Dnmt1^ΔPrx1^* chondrocytes (Figures 3C, D). These *in vitro* results support the *in vivo* findings that Dnmt1 regulates both cell proliferation and differentiation. To verify chondrocyte differentiation at the cellular level, real-time RT-PCR was performed using primary cultured chondrocytes treated with or without BMP2 stimulation. In *Dnmt1^ΔPrx1^* chondrocytes, gene expression levels of *Col2a1*, a marker of early chondrocyte differentiation, were significantly decreased, and gene expression of late markers such as *Runt-related transcription factor 2* (*Runx2*) and *Matrix metalloproteinase-13* (*Mmp13*) was significantly increased before BMP2 treatment. Genes related to osteoblast differentiation and calcification, such as *Osterix* (*Sp7*), *Alkaline phosphatase* (*Alp*), *Osteopontin* (*Opn*), and *Osteocalcin* (*Ocn*), were also significantly increased in *Dnmt1^ΔPrx1^*chondrocytes (Figures 3E). Since cartilage calcification is known to be accompanied by apoptosis, we performed TUNEL staining ^12, 13, 14^. As previously reported, hypercalcification in *Dnmt1^ΔPrx1^* chondrocytes was accompanied by an increased population of apoptotic cells (Figure S3F, G). These data suggested that Dnmt1 deficiency in primary cultured chondrocytes was associated with dysregulated proliferation, differentiation, and mineralization, which corresponds to the deficiencies in long bone formation seen for *Dnmt1^ΔPrx1^* mice.

**Figure 3.**
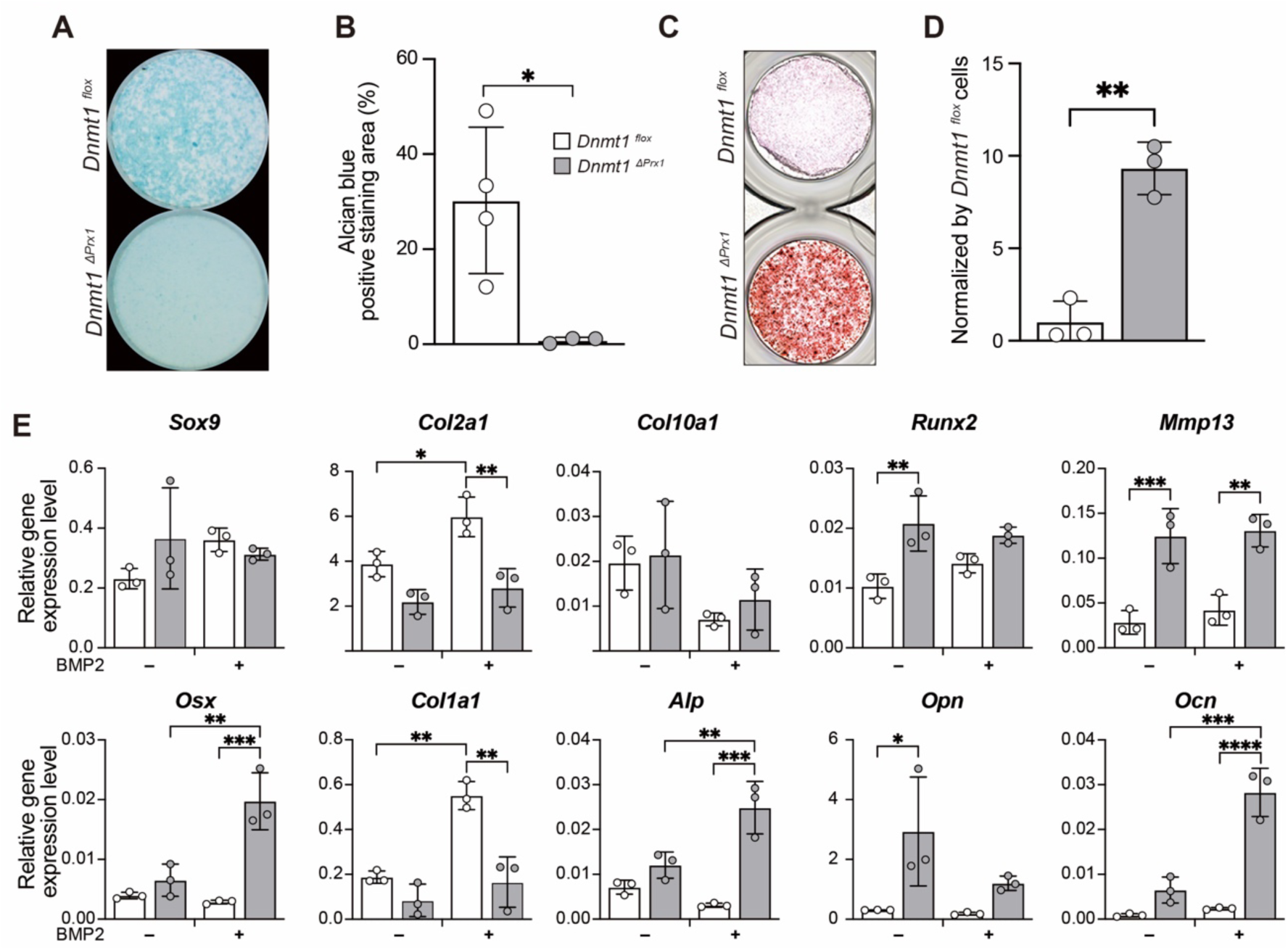
Decreased extracellular matrix production and accelerated mineralization of Dnmt1-deficient chondrocytes. (A) Cartilage matrix synthesis activity in primary cultured chondrocytes obtained from *Dnmt1^flox^* and *Dnmt1^ΔPrx1^* mice as evaluated by Alcian Blue staining. (B) Quantitative analysis of Alcian Blue staining area for *Dnmt1^flox^* (n=4) and *Dnmt1^ΔPrx1^*(n=3) chondrocytes. (C) Mineralization activity of primary cultured chondrocyte obtained from *Dnmt1^flox^* and *Dnmt1^ΔPrx1^*mice as evaluated by Alizarin Red staining. (D) Quantitative analysis of Alizarin Red staining *Dnmt1^flox^* (n=3) and *Dnmt1^ΔPrx1^* (n=3) chondrocytes normalized relative to *Dnmt1^flox^* chondrocytes. (E) Expression levels of genes related to chondrogenesis, mineralization and osteoblast differentiation in *Dnmt1^flox^* (n=3) and *Dnmt1^ΔPrx1^* (n=3) chondrocytes with or without BMP2 treatment. All data are mean ± s.d. * *p* < 0.05, ** *p* < 0.01, *** *p* < 0.001, **** *p* < 0.0001 (unpaired *t* test or Tukey’s post-hoc test).

### Dnmt1 comprehensively regulates gene expression associated with chondrocyte proliferation and differentiation

Next, to identify mechanisms that may cause the observed phenotypes, we comprehensively explored the gene expression profile of Dnmt1-deficient chondrocytes by RNA-Seq using primary cultured chondrocyte treated with or without BMP2. A principal component analysis (PCA) to compare *Dnmt1^flox^*and *Dnmt1^ΔPrx1^* chondrocytes in the presence or absence of BMP2 showed disparate gene expression profiles for cells from *Dnmt1^flox^* and *Dnmt1^ΔPrx1^*mice (Figures S4A, B). Subsequent expression analyses visualized by volcano plots showed upregulated expression of multiple genes in *Dnmt1^ΔPrx1^*chondrocytes compared to *Dnmt1^flox^*, regardless of BMP2 treatment. There is an established relationship between decreased DNA methylation and increased gene expression (Figures S4C, D). To verify that gene expression profiles were altered by Dnmt1 deficiency during chondrocyte differentiation, we compared genes that had significant decreases or increases in expression in the presence of BMP2 treatment for each mouse genotype. BMP2 treatment downregulated the expression levels of 673 and 1,054 genes in *Dnmt1^flox^* and in *Dnmt1^ΔPrx1^* chondrocytes, respectively. Among these genes, the expression of 733 was specifically decreased by Dnmt1 deficiency. Enrichment analysis of these 733 genes revealed an enrichment in genes related to “cell cycle” (Figure 4A). Furthermore, the number of genes that were upregulated in the presence of BMP2 treatment was 2,070 and 1,115 for *Dnmt1^flox^* and *Dnmt1^ΔPrx1^*chondrocytes, respectively. Of these, expression of 612 genes was specifically increased by Dnmt1 deficiency, and enrichment analysis revealed an enrichment in genes involved in “ossification” and “skeletal system development” (Figure 4B). In fact, marker genes for late-stage chondrocyte differentiation including *ADAM metallopeptidase with thrombospondin type 1 motif 4* (*Adamts4*) and *Myocyte enhancer factor 2C* (*Mef2c*) were more highly expressed in *Dnmt1^ΔPrx1^* chondrocytes than in *Dnmt1^flox^* chondrocytes. In addition, *Dnmt1^ΔPrx1^*chondrocytes expressed higher levels of osteoclast-enhancing molecules including *TNF superfamily member 11* (*Tnfsf11*) and *Colony stimulating factor 1* (*Csf1*) (Figure S4E). These results further corroborate the phenotypes observed *in vivo* and *in vitro* and suggest that Dnmt1 comprehensively orchestrates gene expression profiles associated not only with chondrocyte proliferation and differentiation but also with regulation of the microenvironment including osteoclast formation.

**Figure 4.**
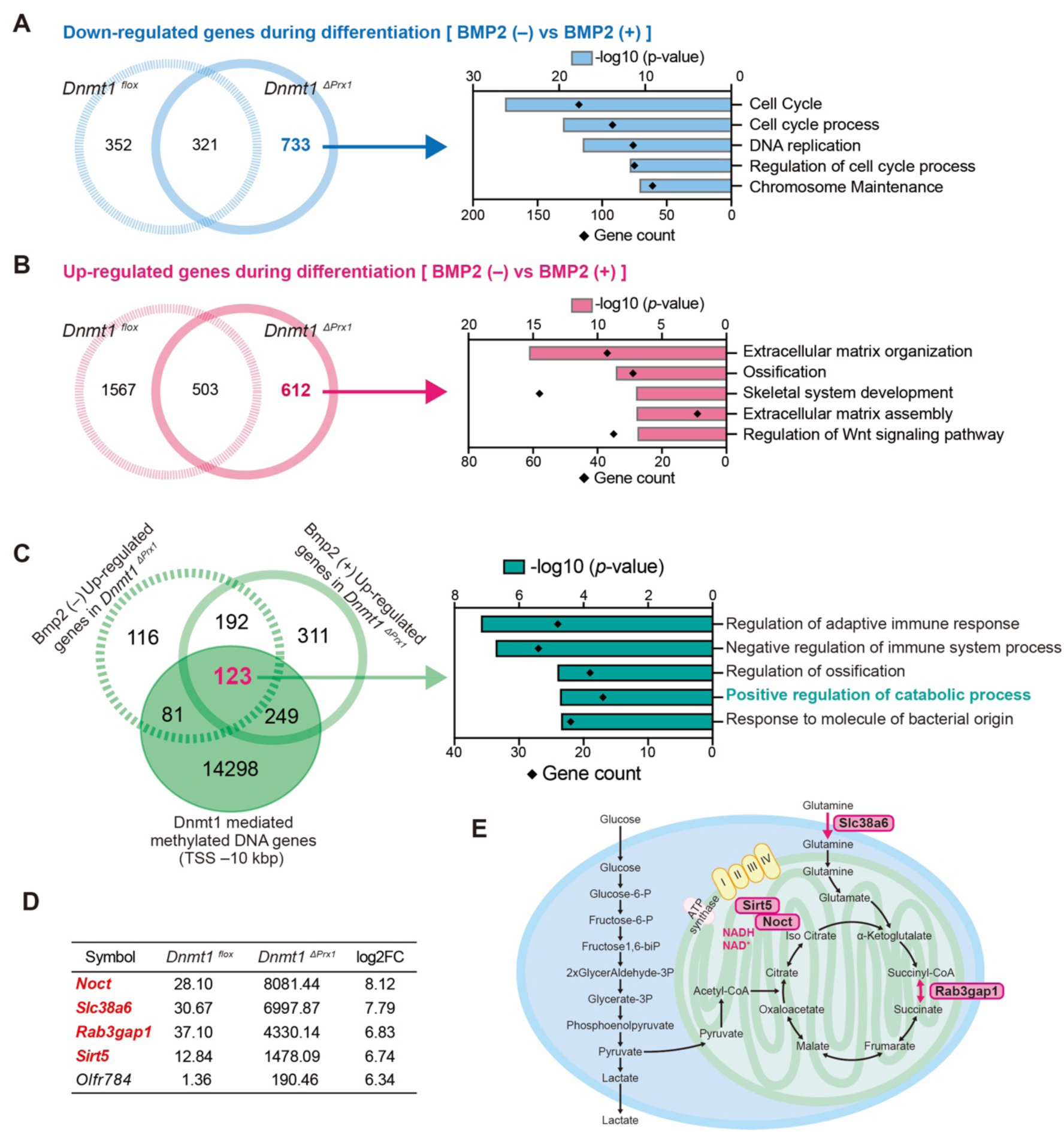
Decreased DNA methylation associated with Dnmt1 deficiency enhances expression of genes related to energy metabolism. (A) Left panel: Venn diagram of genes that had downregulated expression during differentiation of *Dnmt1^flox^* (Light blue dashed circles) and *Dnmt1^ΔPrx1^*(Light blue solid circles) chondrocytes. Right panel: Enrichment analysis of specific downregulated genes (733 genes) during differentiation of *Dnmt1^ΔPrx1^*chondrocytes. (B) Left panel: Venn diagram of genes that were upregulated during differentiation of *Dnmt1^flox^* (Pink dashed circles) and *Dnmt1^ΔPrx1^*(Pink solid circles) chondrocytes. Right panel: Enrichment analysis of *Dnmt1^ΔPrx1^*specific upregulated genes (612 genes) during differentiation of *Dnmt1^ΔPrx1^*chondrocytes. (C) Left panel: Venn diagram to narrow down genes having increased expression accompanying decreases in methylated DNA due to Dnmt1 deficiency. Right panel: Enrichment analysis of genes (n=123) having increased expression with decreases in methylated DNA due to Dnmt1 deficiency. (D) Top 5 genes sorted according to log2 fold-change in RNA-Seq results for chondrocytes without BMP2 treatment. Genes related to energy metabolism are highlighted in red text. (E) Illustration of genes related to energy metabolism.

### Dnmt1 directly regulates expression of genes associated with energy metabolism

To clarify genome-wide changes in DNA methylation status associated with Dnmt1 deficiency and to comprehensively analyze transcriptional regulation by DNA methylation, we next isolated methylated DNA from primary cultured chondrocytes obtained from *Dnmt1^flox^*and *Dnmt1^ΔPrx1^*mice. The methylated DNA was then subjected to next generation sequencing (MBD-seq). The amount of methylated DNA collected using methyl CpG binding domain protein 2 (MBD2) beads was significantly lower for *Dnmt1^ΔPrx1^* chondrocytes than for *Dnmt1^flox^* chondrocytes (Figure S4F) and the peak count frequency of the transcription start site (TSS) region of methylated DNA was also significantly lower in *Dnmt1^ΔPrx1^* chondrocytes (Figure S4G). This result is consistent with the finding that *Dnmt1^ΔPrx1^* chondrocytes had a higher number of upregulated genes than downregulated genes compared to the control *Dnmt1^flox^*chondrocytes (Figure S4C, D). To ascertain the loci of Dnmt1-mediated methylated DNA, we performed MACS2 peak calling using *Dnmt1^flox^* chondrocytes as the ‘treatment’ and *Dnmt1^ΔPrx1^* chondrocytes as the ‘control’. These analyses identified 14,751 gene loci that were targets of Dnmt1-mediated methylation. Combining this result with the RNA-Seq data, we investigated the upregulated genes in *Dnmt1^ΔPrx1^* chondrocytes that overlapped with methylated DNA peaks associated with Dnmt1 activity in chondrocytes that were and were not treated with BMP2 to identify genes that are direct targets of Dnmt1 methylation. We found 123 genes that had Dnmt1-mediated peaks within 10 kb of the TSS (Figure 4C). Gene enrichment analysis revealed that these 123 genes were enriched in the immune system, mineralization, and catabolic process (Figure 4C). Furthermore, 4 of the 5 most highly-ranked genes (Figure 4D), including *Nocturnin* (*Noct*), *Solute carrier family 38 member 6* (*Slc38a6*), *RAB3 GTPase activating protein catalytic subunit 1* (*Rab3gap1*), and *Sirtuin 5* (*Sirt5*), are involved in energy metabolism (Figure 4E) ^15, 16, 17, 18, 19, 20^. Indeed, methylated DNA peaks around the TSS of these genes were reduced in *Dnmt1^ΔPrx1^* chondrocytes (Figure S4H). In support of these data, *Dnmt1^ΔPrx1^* chondrocytes had a significant increase in the expression of genes related to energy metabolism, particularly mitochondria-derived genes (Figure S4E). These results suggest that Dnmt1-mediated methylation regulates the expression of genes involved in energy metabolism and maintains normal chondrocyte function.

### Highly accelerated energy metabolism of *Dnmt1^ΔPrx1^* chondrocytes

We next explored whether the loss of Dnmt1 affects the energy metabolism of chondrocytes. First, glutamine uptake and mitochondrial metabolism were measured to assess the TCA cycle. Glutamine uptake was downregulated in *Dnmt1^ΔPrx1^* chondrocytes compared to *Dnmt1^flox^*chondrocytes (Figure 5A), whereas mitochondrial metabolism was markedly elevated in *Dnmt1^ΔPrx1^*chondrocytes (Figure 5B). Meanwhile, glucose uptake was unaffected by *Dnmt1* knockout (Figure S5A), but glycolysis activity was enhanced (Figure S5B). LC/MS was next carried out to measure changes in metabolites associated with bioenergetic pathways. Levels of almost all major energy metabolites involved in both the TCA cycle and glycolysis, including coenzymes, were increased in *Dnmt1^ΔPrx1^* chondrocytes compared to *Dnmt1^flox^* chondrocytes (Figure 5C, S5C). Among them, levels of S-adenosyl-methionine (SAM) and 2-ketoglutarate (2KG) were higher in *Dnmt1^ΔPrx1^* chondrocytes than in control *Dnmt1^flox^* chondrocytes. SAM is a well-known substrate for DNA methylation, while 2KG is a substrate for demethylation of methyl groups. The higher levels of SAM and 2KG in *Dnmt1^ΔPrx1^* chondrocytes suggest that genome-wide reductions in DNA methylation level due to Dnmt1 deficiency decreased demand for DNA demethylation, and were accompanied by increases in SAM and 2KG in *Dnmt1^ΔPrx1^*chondrocytes. In particular, 2KG is important for the TCA cycle, and is a representative energy metabolite that had increased levels in *Dnmt1^ΔPrx1^*chondrocytes.

**Figure 5.**
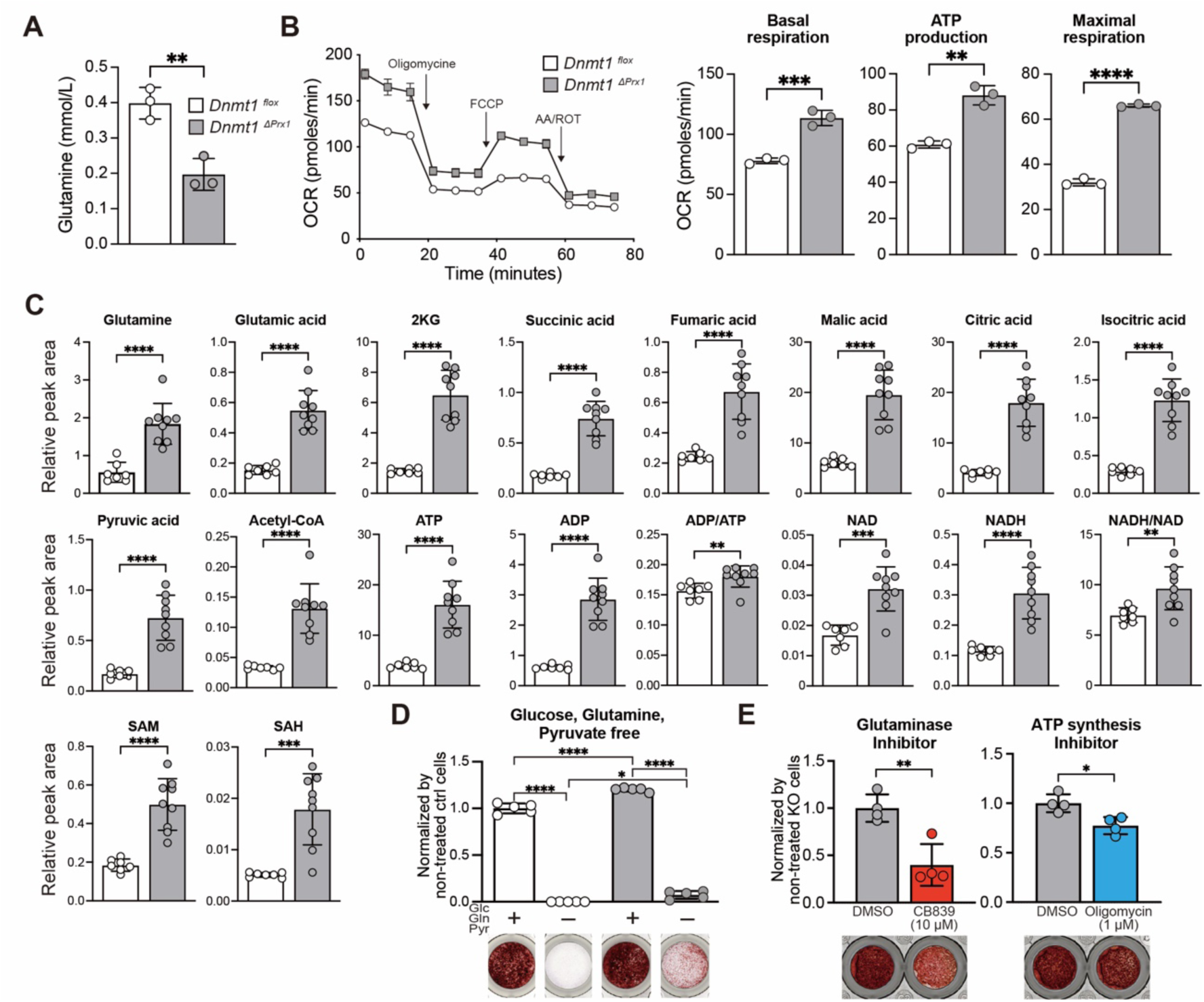
Increased energy metabolism and promotion of mineralization in Dnmt1-deficient chondrocytes. (A) Glutamine uptake in *Dnmt1^flox^* (n=3) and *Dnmt1^ΔPrx1^* (n=3) chondrocytes. (B) Left panel: Oxygen consumption rate (OCR) assessed after addition of oligomycin, carbonyl cyanide 4-(trifluoromethoxy) phenylhydrazone (FCCP), and antimycin A/rotenone (AA/ROT) at the indicated times. Right panel: Basal respiration, ATP production and maximal respiration measured for *Dnmt1^flox^* (n=3) and *Dnmt1^ΔPrx1^*(n=3) chondrocytes. (C) Relative quantification of energy metabolites by metabolome analysis of *Dnmt1^flox^* (n=7) and *Dnmt1^ΔPrx1^* (n=9) chondrocytes. (D) Calcification activity in primary cultured chondrocytes in the presence or absence of glucose, glutamine, and pyruvate as evaluated by Alizarin Red staining of *Dnmt1^flox^* (n=5) and *Dnmt1^ΔPrx1^* (n=5) chondrocytes. (E) Calcification activity in *Dnmt1^ΔPrx1^* chondrocytes in the presence of energy metabolism inhibitors (CB839: Glutaminase inhibitor (n=4), Oligomycin: ATP synthesis inhibitor (n=4)) as evaluated by Alizarin Red staining. All data are mean ± s.d. * *p* < 0.05, ** *p* < 0.01, *** *p* < 0.001, **** *p* < 0.0001 (unpaired *t* test or Tukey’s post-hoc test).

Enhanced energy metabolism is required for cell differentiation and mineralization ^21, 22, 23, 24^. Here we examined whether enhanced mineralization seen in *Dnmt1^ΔPrx1^* chondrocytes can be suppressed by removing glucose, glutamine, and pyruvate from the culture medium. As expected, essentially no calcification in *Dnmt1^flox^* chondrocytes was observed in the absence of glucose, glutamine, and pyruvate, but slight calcification levels did persist in *Dnmt1^ΔPrx1^* chondrocytes (Figure 5D), suggesting that aberrant energy metabolism pathways may be engaged in *Dnmt1^ΔPrx1^* chondrocytes. Furthermore, treatment with TCA cycle inhibitors, such as glutaminase (CB839) or ATP synthesis (Oligomycin) inhibitors, suppressed the abnormally enhanced energy metabolism (Figure S5D) and mineralization (Figure 5E) seen in *Dnmt1^ΔPrx1^* chondrocytes. These results support a role for Dnmt1 activity at both a transcriptional and metabolic level that regulates chondrocyte energy metabolism and in turn cartilage calcification.

### DNMT1 deficiency accelerates energy metabolism and differentiation in human chondrocytes

Finally, to determine whether the mechanisms defined in mice could be applicable to humans, we performed similar analyses using human chondrocytes. Human chondrocytes used in these studies were isolated from knee articular cartilage that was removed and discarded during surgical procedures. First, the gene expression levels were suppressed by siRNA against *DNMT1*, and the knockdown efficiency was confirmed by RT-qPCR and immunostaining. DNMT1 gene and protein expression levels were decreased in chondrocytes treated with two types of siRNA against *DNMT1*compared to chondrocytes treated with siControl (Figure S6A-C). On the other hand, gene expression levels of UHRF1 did not differ between siDNMT1- and siControl-treated chondrocytes (Figure S6A). Gene expression levels of *NOCT*, *SIRT5* and *SLC38A6* were high in *Dnmt1^ΔPrx1^* chondrocytes and upregulated in siDNMT1-treated chondrocytes (Figure 6A). Finally, cell proliferation, differentiation and energy metabolism of human chondrocytes treated with siDNMT1 were analyzed. The cell proliferation rate of chondrocytes was reduced by siDNMT1 treatment (Figure S6D). Expression of marker genes for the early stage of chondrocyte differentiation including *SOX9*, *COL2A1* and *ACAN* was downregulated whereas expression of marker genes for the late stage of chondrocyte differentiation, such as *RUNX2, COL10A1* and *MMP13*, was upregulated in siDNMT1 chondrocytes compared to siControl chondrocytes (Figure 6B). Genes related to ossification such as *COL1A1*, *ALPL* and *BGLAP* were also upregulated in siDNMT1-treated chondrocytes. Furthermore, mitochondrial activity in siDNMT1-treated chondrocytes was significantly increased compared to siControl chondrocytes (Figure 6C). These results demonstrated that DNMT1-deficient human chondrocytes exhibited similar characteristics to those of *Dnmt1^ΔPrx1^* chondrocytes and strongly suggest that DNMT1 plays essential roles in both proliferation and differentiation of chondrocytes through the regulation of energy metabolism.

**Figure 6.**
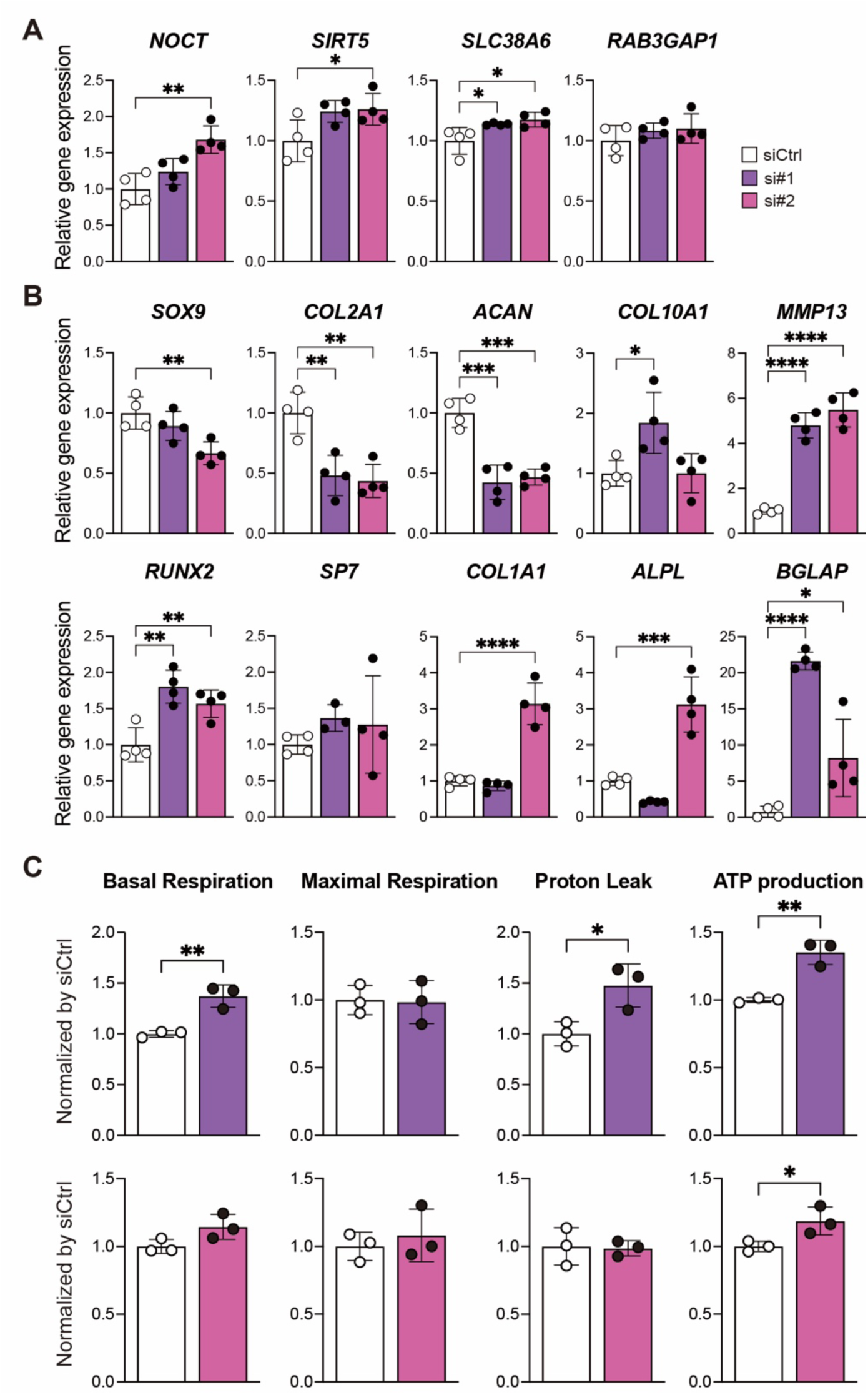
Accelerated ATP production and differentiation in human chondrocytes with DNMT1 deficiency. Gene expression levels of (A) genes related to energy metabolism related and (B) markers of chondrogenesis differentiation after differentiation stimulus by BMP2 as determined by real-time RT-PCR (n=4). (C) Basal respiration, maximal respiration, proton leak and ATP production calculated from OCR (n=3). (Upper panel: siCtrl vs. siDNMT1 #1, Lower panel: siCtrl vs. siDNMT1 #2) All data are mean ± s.d. * *p* < 0.05, ** *p* < 0.01 (unpaired *t* test or Tukey’s post-hoc test).

## Discussion

In this study, Dnmt1 deficiency in limb mesenchymal cells was associated with limb shortening and enhancement of ossification (Figure 1B-E, S1A-C, 2E). Dnmt1 plays a crucial role in the maintenance of DNA methylation, one of the most well-known epigenetic modifications. DNA methylation induces structural changes in chromatin and accumulation of methylation that in turn contributes to regulation of transcription ^3^. Establishment of unique gene expression patterns during cell proliferation and differentiation is attributed to DNA methylation, and inhibition of DNA methylation leads to abnormal development ^25, 26^.

Here, RNA-Seq analysis of Dnmt1-deficient chondrocytes revealed osteoblastic phenotypes (Figure 4A, B, S4E) that were consistent with previous reports, indicating that DNMT1 activity affects osteoblast differentiation that negatively regulates mineralization ^10, 11^. In addition, Dnmt1 expression decreases concurrent with chondrocyte differentiation (Figure 2A, S3A). Taken together with the observed phenotype of postnatal arrest of long bone growth in Dnmt1-deficient mice, we conclude that Dnmt1 helps regulate longitudinal growth of long bones in part by controlling mineralization of postnatal growth plate chondrocytes. *Dnmt1^ΔPrx1^* mice also exhibited greater shortening of bone length compared to mice with limb mesenchymal cell-specific *Uhrf1* knockout ^5^. This difference could indicate that mechanisms for maintenance of DNA methylation involving Dnmt1 have a larger role in limb development than the multi-functional Uhrf1. However, the mechanism by which Dnmt1 regulates mineralization in chondrocytes remained unclear.

To elucidate the detailed molecular mechanism of Dnmt1 in chondrocyte mineralization here we carried out an integrated RNA-Seq and MBD-Seq analysis. Genome-wide integrated analysis indicated that Dnmt1 regulated not only expression of genes involved in ossification, but also that of genes related to energy metabolism (Figure 4C-E). Indeed, calcification was enhanced (Figure 3C, D), and also levels of nearly all major energy metabolites, including ATP, required for mineralization, as well as SAM and 2KG, associated with DNA methylation, were increased in *Dnmt1^ΔPrx1^* chondrocytes (Figure 5C). These results were similar to those for Dnmt1-deficient human articular chondrocytes in which expression of genes related to energy metabolism and ATP production was upregulated (Figure 6C). ATP is essential for cell survival, proliferation, and differentiation. In chondrocytes in particular, ATP is a substrate for pyrophosphate (PPi) feeding and inorganic phosphate (Pi) synthesis to produce hydroxyapatite ^27^. Previously, transgenic mice overexpressing *Ankylosis* (*Ank*), a key regulator of physiological mineralization that regulates the Pi/PPi balance, had shortened spines and enhanced mineralization ^28^. A primary molecular mechanism for ATP production is the TCA cycle, which involves both 2KG and SAM. During the TCA cycle, SAM is converted to SAH, which eventually leads to the formation of succinyl CoA and pyruvate that further feed back into the cycle ^29^. 2KG also functions as a substrate for DNA demethylation mediated by TET family proteins, which promote DNA demethylation by converting 5-methylcytosine (5mC) to 5-hydroxymethylcytosine (5hmC) ^4^. In *Dnmt1^ΔPrx1^* chondrocytes, DNA methylation is reduced genome-wide and concurrently the demand for DNA demethylation is reduced leading to accumulation of 2KG. This accumulation of 2KG decreases collagen synthesis via upregulation of collagen prolyl hydroxylases and enhances ossification that leads to shortening of long bones ^24^. Previous reports indicated that proper amounts of TCA cycle intermediates are vital for the development and homeostasis of cartilage and bone ^22^.

Meanwhile, mitochondrial transcription factor A (Tfam), a nuclear DNA-encoded protein, is needed to maintain mtDNA. Disruption of *Tfam* in chondrocytes causes severe ATP depletion via decreased mtDNA that results in suppressed chondrocyte mineralization with long bone shortening ^30^. Mice deficient in both *Tfam*/*Hif1a* had improved chondrocyte mineralization and bone length through upregulated production of metabolites involved in the TCA cycle ^13^. Abnormal glutamine metabolism associated with the Hif1a-induced 2KG stabilization pathway promotes mineralization accompanied by increased apoptosis that impairs bone growth ^23, 31, 32^. However, the mechanisms associated with regulation and control of energy metabolism and energy supply in cartilage, an avascular tissue, during chondrocyte differentiation were not clear. In this study, we discovered a novel comprehensive intracellular energy metabolism regulation mechanism, in which Dnmt1-mediated DNA methylation controls energy metabolism by regulating expression of genes related to energy metabolism and restricting intracellular consumption of SAM and 2KG associated with DNA demethylation.

Increased expression of Dnmt1 has been associated with the progression of OA ^9, 33^ while abnormalities related to energy metabolism are involved in OA onset ^34^. Questions concerning the role of Dnmt1 activity in human and mouse chondrocytes remain to be addressed, such as how Dnmt1 expression is regulated during differentiation and whether Dnmt1-mediated changes in energy metabolism are relevant to other age-related pathologies, including OA.

In conclusion, our findings suggest that Dnmt1-mediated maintenance of appropriate DNA methylation during differentiation governs chondrocyte differentiation/mineralization by altering energy metabolism through the regulation of expression of genes related to chondrocyte differentiation and of genes related to the supply of metabolites that in turn control bone elongation.

## Methods

### Animals

*Dnmt1 flox* mice were kindly provided by Prof. Rudolf Jaenisch. *Dnmt1* flox mice were crossed with *Prrx1*-Cre mice [B6.Cg-Tg(Prrx1-cre)1Cjt/J, Jackson Laboratory] to generate *Prrx1*-Cre; *Dnmt1^flox/flox^*(*Dnmt1^Δprx1^*) mice. Littermate *Dnmt1 flox* (*Dnmt1^flox^*) mice were used as a control. To identify molecules regulated by Dnmt1 that are not dependent on sex, both male and female mice were used. All mice were housed in a specific pathogen-free facility under climate-controlled conditions with a 12-h light/dark cycle and were provided with water and standard diet (MF, Oriental Yeast, Japan) *ad libitum*. All animals were maintained and used according to the experimental protocol approved by the Animal Experiment Committee in Ehime University, Japan.

### Skeletal preparation

Neonates were skinned, eviscerated, and fixed in 100% ethanol for 24 hours before transfer to 100% acetone for 24 hours. Skeletal samples were stained with 0.015% Alcian Blue followed by 0.005% Alizarin Red and 0.05% glacial acid dissolved in 70% ethanol for 4-7 days before sequential clearance in 1% KOH with glycerol. Cartilage and mineralized bone were stained blue and red, respectively.

### Bone length calculation and images

Long bones were harvested and fixed with 4% paraformaldehyde (PFA) overnight at 4 °C and soaked in 70% ethanol. Bone length was calculated using ImageJ from images taken with a soft X-ray device (Sofron) ^35, 36^. A Scanco Medical μCT35 system (SCANCO Medical) was used for microcomputed tomography (µCT).

### Histological analysis

Mice were euthanized and the tibiae and femurs were harvested. The collected tissue was fixed with 4% PFA overnight at 4 °C, followed by decalcification with EDTA. For *in vivo* BrdU uptake assays, 100 mg/kg BrdU (Abcam: ab142567) was injected intraperitoneally 24 hours before sampling. Long bone samples were embedded in paraffin after dehydration and 5-7 μm-thick paraffin sections were cut with a microtome (RM2255, Leica Biosystems). Sections were stained with von Kossa, Alcian Blue and also used for immunohistochemistry (IHC). For IHC, deparaffinized sections were incubated in 2.5% hyaluronidase (Sigma-Aldrich) solution for 30 min at room temperature before boiling for 45 min at 90 °C in 0.05% citraconic acid solution (ImmunoSaver; Wako) to retrieve antigens. After blocking for 60 minutes at room temperature in PBS containing 1% bovine serum albumin (BSA) and 0.02 % Triton, the sections were incubated overnight at 4 °C with primary antibodies against Dnmt1 (1:50, D63A6, CST), Uhrf1 (1:50, sc-373750, Santa Cruz), BrdU (1:250, ab6326, abcam), Sp7 (1:1000, ab209484, abcam) or Ctsk (1:50, sc-48353, Santa Cruz). The sections were washed again with PBS and signals were visualized after incubating with secondary antibodies for 60 min at room temperature followed by addition of TrueVIEW (SP-8400, Vector Laboratories) to suppress autofluorescence. The area of proliferative and hypertrophic cartilage was measured using ImageJ according to the cell morphology in regions of interest (ROI) in the tibia.

### Cell culture

For isolation of primary chondrocytes, knee cartilage was harvested from postnatal day 3-5 mice and treated with 3 mg/ml collagenase D (11088866001, Roche) in DMEM (10569010, Gibco) supplemented with 1% antibiotic-antimycotic solution (15240062, Gibco) for 90 minutes. The connective tissue was then thoroughly removed by washing and the tissue was incubated overnight with fresh medium containing collagenase. Cells were cultured in DMEM supplemented with 10% fetal bovine serum (FBS, Nichirei Biosciences) and 1% antibiotic/antimycotic solution (15240062, Gibco) at 37 °C and 5% CO_2_. To induce differentiation, cells were seeded at confluence, cultured with 100 ng/ml BMP2 (CK-B2, Osteopharma), 50 μg/ml ascorbic acid and 10 mM β-glycerophosphate, and stained with Alcian Blue and Alizarin Red. To inhibit energy metabolism, cells were incubated with 10 μM CB839 (glutaminase inhibitor, Selleckchem) and 1 μM oligomycin (ATP synthesis inhibitor, Adipogen).

### Immunocytochemistry

Primary cultured chondrocytes were seeded onto chamber slides (177445PK, Lab-Tek) at a density of 1×10^4^ cells per well. For BrdU incorporation assays, the medium was replaced with medium containing 10 μM BrdU. The cells were fixed with 4% PFA for 5 minutes at room temperature and then permeabilized with 0.5% Triton in PBS for 10 minutes. Non-specific binding was blocked by incubating cells with 1% BSA and 0.02% Triton in PBS for 1 hour at room temperature. Cells were then incubated overnight at 4 °C with primary antibodies against target proteins (Dnmt1, BrdU) that were diluted in blocking buffer. After washing with PBS, cells were incubated with fluorophore-conjugated secondary antibodies for 1 hour at room temperature in the dark. Nuclei were counterstained with 4’,6-diamidino-2-phenylindole (DAPI) for 5 minutes. Finally, coverslips were mounted onto glass slides using mounting medium and sealed with nail polish. Immunofluorescence images were acquired using a fluorescence microscope.

### Real time RT-PCR

Total RNA was extracted using Isogene (Nippon Gene) and an RNeasy spin column kit (QIAGEN), and then treated with DNase I (QIAGEN). cDNA was synthesized from the total RNA using PrimeScript RT Master Mix (Takara Bio) and real-time RT-PCR performed using TB-Green Premix Ex Taq II (Takara Bio) with Thermal Cycler Dice (Takara Bio) according to the manufacturer’s instructions. Gene expression levels were normalized using the housekeeping gene Gapdh. Primer sequences are listed in the Supplementary Table 1.

### RNA-Seq analysis

Primary cultured chondrocytes that were and were not treated with BMP2 for 3 days were used for RNA-Seq. Total RNA was extracted using an RNeasy spin column kit and the quality verified using an Agilent 2100 Bioanalyzer. RNA-Seq analysis was performed as previously described ^5, 37^. RNA-seq libraries were prepared using an NEBNext Ultra II Directional RNA Library Prep Kit according to the manufacturer’s instructions and subsequently validated for an average size of about 370 bp using a 2100 Bioanalyzer and an Agilent DNA1000 kit. Sequencing of paired-end reads (150 bp) was performed with a NovaSeq 6000 S4 Reagent Kit on a NovaSeq 6000 (Illumina). Sequence data were analyzed by RaNA-Seq ^38^. The RNA-Seq data set was deposited in GEO under accession number GSE270640.

### MBD-Seq analysis

MBD-Seq was performed for genome-wide analysis of methylated DNA regions as previously described ^5, 37^. Methylated DNA was enriched by MBD2-mediated precipitation and subjected to next-generation sequencing as follows. Extracted DNA from chondrocytes was sonicated with a Covaris sonicator (M&S Instruments) to obtain approximately 300bp fragments. MBD2-mediated enrichment of methylated DNA was performed using an EpiXplore Methylated DNA Enrichment Kit (Takara Bio) according to the manufacturer’s instructions. The amount of enriched methylated DNA from 1 μg total DNA was measured using a Quantus Fluorometer (Promega). Libraries for MBD-Seq analysis were prepared using a NEBNext Ultra II DNA Library Prep Kit for Illumina according to the manufacturer’s instructions and validated for an average size of about 550-600 bp using a High Sensitivity DNA Assay kit. Sequencing of paired-end reads (150 bp) was performed using a NovaSeq 6000 S4 Reagent Kit with a NovaSeq 6000 (Illumina) and mapped on the mouse genome (mm10) using Galaxy ^39^. The MBD-Seq data set was deposited in GEO under accession number GSE270641.

### Analysis of sequencing data

Using RNA-Seq results, differentially expressed genes that exhibited more than two-fold increased or decreased expression levels in *Dnmt1^ΔPrx1^* mice relative to *Dnmt1^flox^*mice were extracted. Gene Ontology analyses were performed on the extracted genes using MetaScape ^40^. For MBD-Seq, peak calling was performed by MACS2 and integrative analyses were carried out using Galaxy ^39^.

### Glucose and glutamine uptake assay

Primary cultured chondrocytes were seeded at 2 × 10^4^ cells/well in 96-well plates the day before measurement. Glucose and glutamine concentrations in the medium were measured using a Glucose Assay Kit-WST (DOJINDO, Japan) and Glutamine Assay Kit-WST (DOJINDO, Japan), respectively, and uptake was calculated according to the manufacturer’s instructions. Absorbance at 450 nm was measured using a Multiskan SkyHigh Microplate Spectrophotometer (Thermo Fisher).

### Cellular metabolism assay

Metabolic flux was measured using a Seahorse XFp Flux Analyzer (Seahorse Bioscience). Before initiating the assay, primary cultured chondrocytes were seeded at a density of 3 × 10^4^ cells per well into an 8-well Seahorse culture plate (Seahorse Bioscience). For OCR analysis, cells were cultured for 1 hour in DMEM supplemented with 10 mM glucose, 1 mM pyruvate and 2 mM glutamine (Seahorse Bioscience) and equilibrated at 37 °C in a CO_2_-free atmosphere. Following three basal measurements, 2 μM oligomycin, 0.25 μM carbonyl cyanide 4-(trifluoromethoxy) phenylhydrazone (FCCP) and 0.5 μM antimycin A/rotenone were sequentially injected into the plate. For analysis of the ECAR, cells were cultured for 1 hour in DMEM supplemented with 2 mM glutamine and equilibrated at 37 °C in a CO_2_-free atmosphere. After two basal measurements, 10 mM glucose and 2 μM oligomycin were sequentially injected. Analysis data was calculated using Wave Desktop software (Seahorse Bioscience).

### Metabolome analysis by mass spectrometry

For metabolome analysis, primary cultured chondrocytes were seeded in culture dishes without BMP2 treatment. The cells were washed with cold PBS, quenched with cold methanol, and collected by scraping. Sample preparation for mass spectrometry was performed as previously reported ^41^. Anionic polar metabolites were analyzed via ion chromatography (Dionex ICS-5000+ HPIC system, Thermo Fisher Scientific) with a Dionex IonPac AG11-HC-4 μm guard column (2 mm i.d × 50 mm, 4 μm particle size, Thermo Fisher Scientific) and a Dionex IonPac AS11-HC-4 μm column (2 mm i.d × 250 mm, 4 μm particle size, Thermo Fisher Scientific) coupled to a Q Exactive mass spectrometer (Thermo Fisher Scientific) (IC/MS). Cationic polar metabolites were analyzed via liquid chromatography (LC) (Dionex UltiMateTM 3000 UHPLC System, Thermo Fisher Scientific) with a Discovery HS F5 column (2.1 mm i.d × 150 mm, 3 μm particle size, Sigma-Aldrich) coupled to a Q Exactive instrument (PFPP-LC/MS). Acyl-CoAs and acyl-carnitines were analyzed using LC (Nexera X2 UHPLC system, Shimadzu Co. Kyoto, Japan) with an Inert Sustain C18 column (2.1 mm i.d × 150 mm, 3 μm particle size, GL Sciences Inc., Tokyo, Japan) coupled to a Q Exactive instrument (C18-LC/MS). Hydrophilic metabolite data from the three platforms were analyzed using Reifycs Cascade software (ver. 1.1.2).

### Isolation of human chondrocytes

Human articular specimens were obtained from patients with OA who underwent knee joint replacement surgery at the Ehime University Hospital. To obtain chondrocytes, articular cartilage tissues were minced and treated with 0.1% trypsin solution and incubated for 30 min at 37 °C with shaking. The connective tissue was then thoroughly removed by washing, followed by treatment with fresh medium containing 3 mg/ml collagenase D for 2-3 hour at 37 °C with shaking. The supernatant was discarded carefully and fresh medium containing 3 mg/ml collagenase D was added before incubation for overnight at 37 °C. Isolated cells were filtered with a 40 μm cell strainer (Falcon) and seeded in culture dishes at 37 °C in a humidified atmosphere of 5% CO_2_. Experiments involving human samples were approved by the IRB of Ehime University. All patients provided informed written consent to participate in the study.

### siRNA experiments using human chondrocytes

siRNA for DNMT1 and scrambled siRNA for as a negative control were purchased from Thermo Fisher Scientific. siRNAs were transfected into human chondrocytes obtained from articular cartilage using Lipofectamine™ RNAiMAX Transfection Reagent (Thermo Fisher Scientific). After treatment with siRNA for 1 week, cells were used in cell counting assays, cellular metabolism assays, immunocytochemistry, and measurement of chondrocyte differentiation marker genes. In addition to the siRNA period described above, total RNA was extracted after an additional 3 weeks of treatment with siRNA and 100 ng/ml BMP2. For the cell counting assay, the same number of human chondrocytes was seeded before siRNA treatment, and 1 week after siRNA treatment, when the cell number was counted and the proliferation rate was calculated.

### Statistical analysis

We used two-tailed unpaired Student’s t tests to analyze differences between two groups with GraphPad Prism 10. ANOVA followed by post-hoc Tukey’s test as appropriate was applied with GraphPad Prism 10. For all graphs, data are represented as the mean ± SD. Statistical significance was accepted when *P* values were less than 0.05. The sample sizes were determined empirically based on our previous experience and review of similar experiments in the literature. The number of animals utilized is detailed in the corresponding figure legends.

## Acknowledgments

We thank the staff at the Division of Medical Research Support, the Advanced Research Support Center (ADRES) and the members of the Division of Integrative Pathophysiology, Proteo-Science Center (PROS), Ehime University, for their technical assistance and helpful support. This study was supported in part by the Japan Society for the Promotion of Science (JSPS) KAKENHI Grant Number JP20K18066 (to YY) as well as JP23689066, JP15H04961, JP19H03786, and JP22H03203 (to YI).

This study was also supported in part by a grant from the Research Support Project for Life Science and Drug Discovery (Basis for Supporting Innovative Drug Discovery and Life Science Research (BINDS)) from AMED (Grant Number JP24ama121055). This work was performed as part of the MEXT Cooperative Research Project Program, Medical Research Center Initiative for High Depth Omics, and CURE:JPMXP1323015486 for the Medical Institute of Bioregulation, Kyushu University.

## Author contributions

YY and Y Imai planned the study and designed the experiments. MT, Y Izumi, and TB performed mass spectrometry analyses, and TK and MT collected and provided human samples. YY performed all experiments and subsequent analyses with support and advice from the co-authors. YY and Y Imai wrote the manuscript with input from the co-authors. In particular, TB provided valuable advice regarding mass spectrometry analyses while MT provided valuable advice regarding experiments involving human samples.

