## Supplemental information for "Dnmt1 determines bone length by regulating energy metabolism of growth plate chondrocytes"

### Figure 1 - figure supplement 1

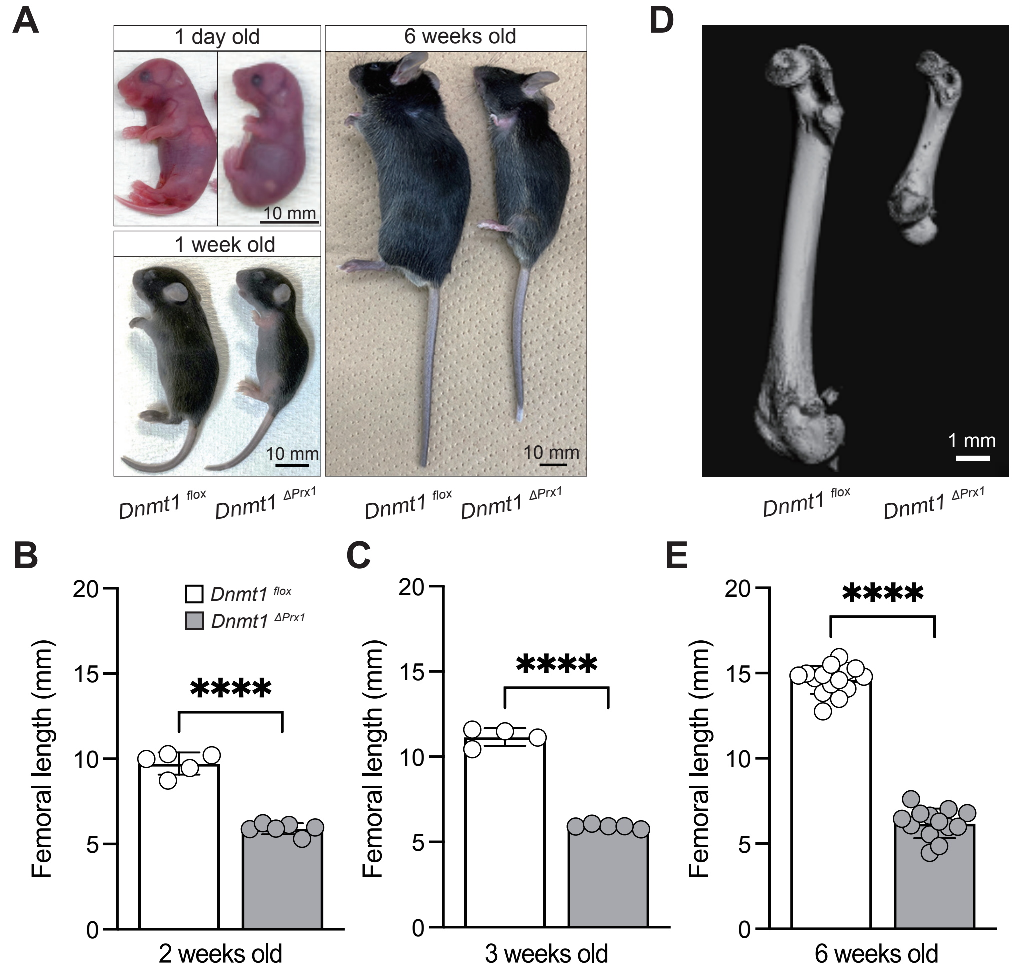


**Figure S1. Long bones of *Dnmt1$^{\Delta Prx1}$* mice show little elongation after birth.**
(A) Representative *Dnmt1$^{flox}$* and *Dnmt1$^{\Delta Prx1}$* mice at different ages (Upper left; 1 day old, lower left; 1 week, right; 6 weeks old). Length of femurs from (B) 2-week-old (*Dnmt1$^{flox}$* (n=5) and *Dnmt1$^{\Delta Prx1}$* (n=6)) and (C) 3-week-old (*Dnmt1$^{flox}$* (n=4) and *Dnmt1$^{\Delta Prx1}$* (n=5)) mice. (D) µCT images of femurs from 6-week-old *Dnmt1$^{flox}$* and *Dnmt1$^{\Delta Prx1}$* mice. (E) Length of femurs from 6-week-old *Dnmt1$^{flox}$* (n=15) and *Dnmt1$^{\Delta Prx1}$* (n=13) mice. All data are mean ± s.d. **** $p < 0.0001$ (unpaired *t* test).

### Figure 2 - figure supplement 2

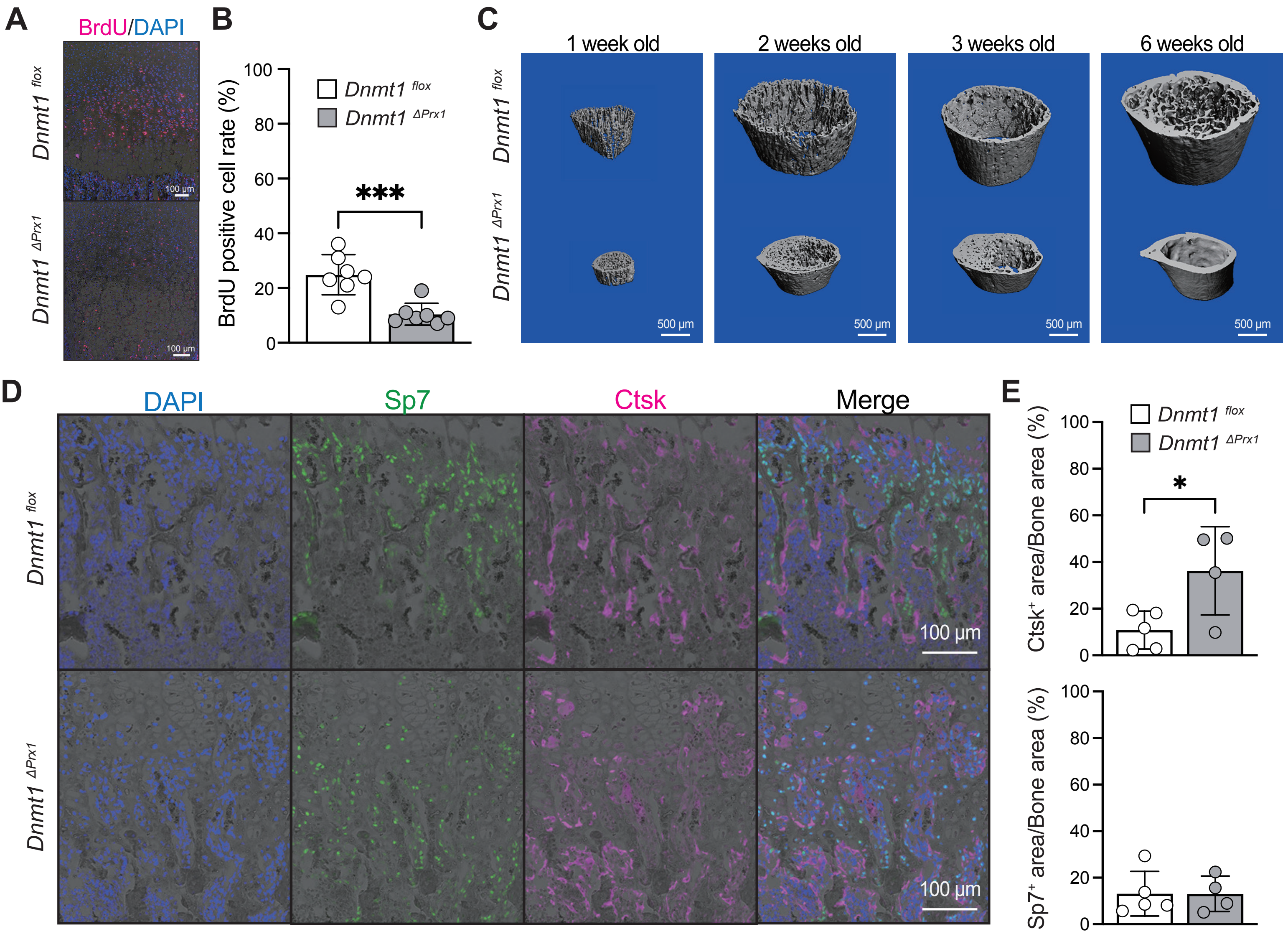


**Figure S2. Cancellous bone in *Dnmt1*$^{\Delta Prx1}$ mice forms early and disappears with age.**
(A) Immunohistochemistry to detect BrdU in proximal tibiae from 1-week-old mice. (B) Quantification of BrdU-positive cells in proximal tibiae from 1-week-old *Dnmt1*$^{flox}$ (n=7) and *Dnmt1*$^{\Delta Prx1}$ (n=7) mice. (C) µCT images of tibial growth plates from 1-, 2-, 3- and 6-week-old mice. Representative data from at least three individual mice are shown. (D) Sp7 (Green), Ctsk (Magenta) and DAPI (Blue) immunohistochemistry of proximal tibiae from 2-week-old mice. (E) Sp7- and Ctsk-positive area per bone area of proximal tibiae from 2-week-old *Dnmt1*$^{flox}$ (n=5) and *Dnmt1*$^{\Delta Prx1}$ (n=5) mice. Data are mean ± s.d. * $p < 0.05$, *** $p < 0.001$ (unpaired *t* test).

### Figure 3 - figure supplement 3

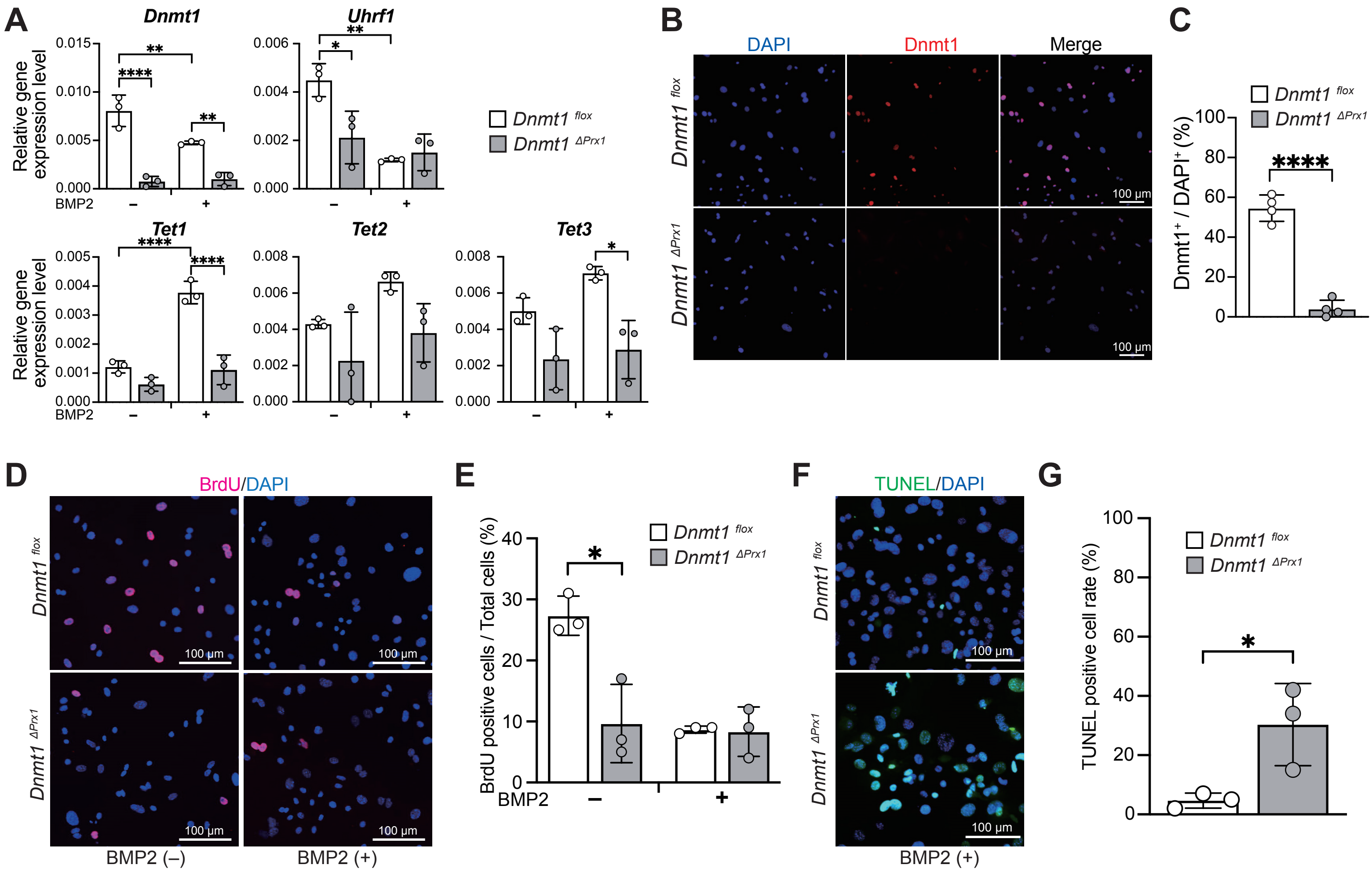


**Figure S3. Reduced cell proliferation and promotion of apoptosis in Dnmt1-deficient chondrocytes.**
(A) Expression levels of genes related to DNA methylation maintenance in *Dnmt1*$^{flox}$ (n=3) and *Dnmt1*$^{\Delta Prx1}$ (n=3) chondrocytes with or without BMP2 treatment. (B) Immunocytochemistry for Dnmt1 (Red) and DAPI (Blue) in primary chondrocyte cultures. (C) Quantification of Dnmt1 expression in *Dnmt1*$^{flox}$ (n=4) and *Dnmt1*$^{\Delta Prx1}$ (n=4) chondrocytes. (D) Immunocytochemistry for BrdU (Red) and DAPI (Blue) in primary chondrocyte cultures with or without BMP2 treatment. (E) Quantitative analysis of BrdU-positive cells in *Dnmt1*$^{flox}$ (n=3) and *Dnmt1*$^{\Delta Prx1}$ (n=3) chondrocytes. (F) TUNEL staining of chondrocytes differentiated by BMP2. (G) Quantitative analysis of TUNEL-positive cells in *Dnmt1*$^{flox}$ (n=3) and *Dnmt1*$^{\Delta Prx1}$ (n=3) chondrocytes. All data are mean ± s.d. * $p < 0.05$, ** $p < 0.01$, **** $p < 0.0001$ (unpaired *t* test or Tukey's post-hoc test).

### Figure 4 - figure supplement 4

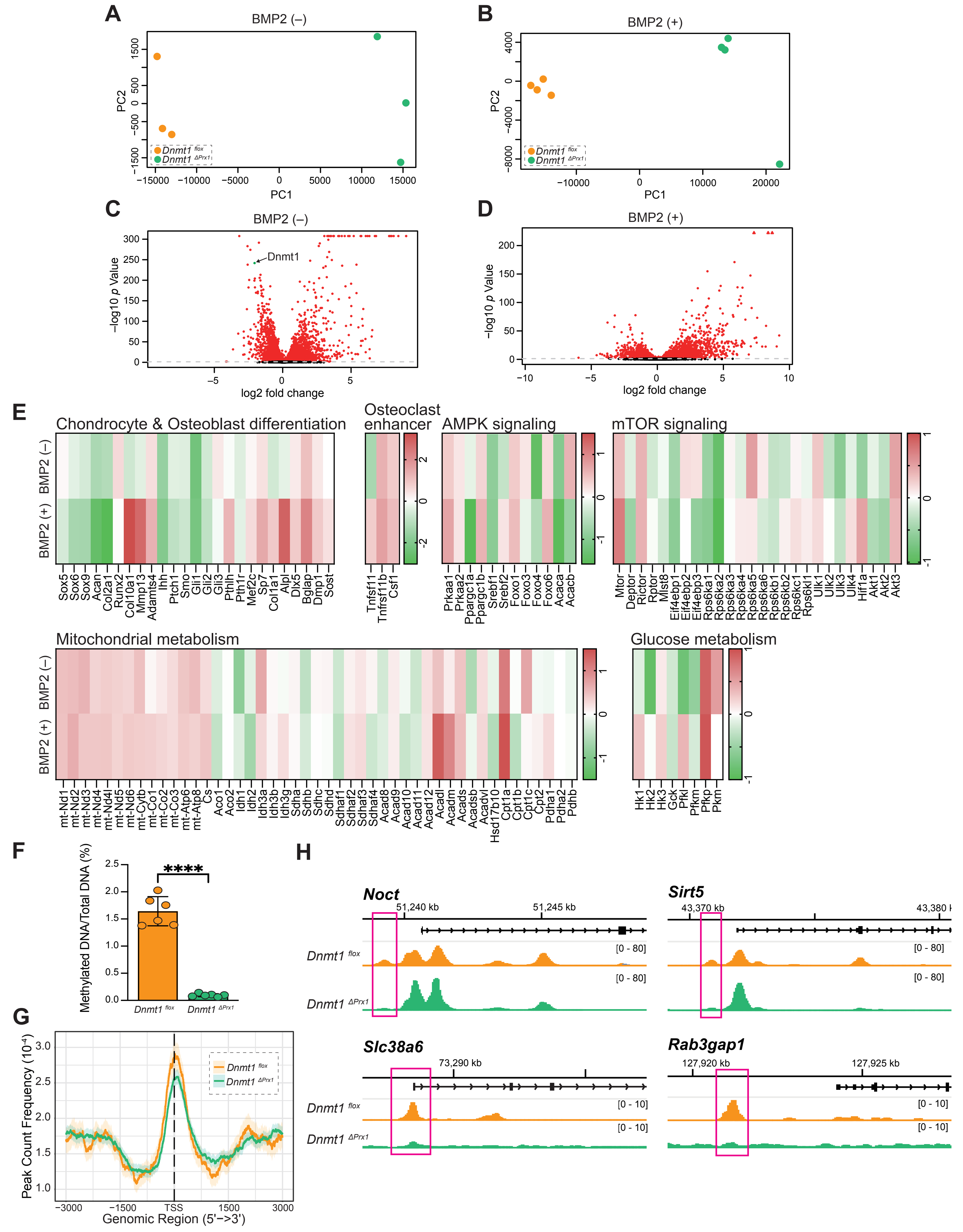

**Figure S4. Decreased DNA methylation in *Dnmt1$^{\Delta Prx1}$* chondrocytes near transcription start sites and gene promoters.**
(A) PCA plot of RNA-Seq of *Dnmt1$^{flox}$* (n=3) and *Dnmt1$^{\Delta Prx1}$* (n=3) chondrocytes without BMP2 treatment. (B) PCA plot of RNA-Seq of *Dnmt1$^{flox}$* (n=4) and *Dnmt1$^{\Delta Prx1}$* (n=4) chondrocytes with BMP2 treatment. (C) Volcano plot for RNA-seq data without BMP2 differentiation stimulus. (D) Volcano plot for RNA-seq data of chondrocytes treated with BMP2. (C, D) Red dots correspond to genes that had significant differential expression (*Padj* < 0.05) in *Dnmt1$^{\Delta Prx1}$* chondrocytes and *Dnmt1$^{flox}$* chondrocytes. (E) Heatmap of genes associated with each term based on RNA-Seq data. Values indicate log2 fold-changes in gene expression levels between *Dnmt1$^{flox}$* and *Dnmt1$^{\Delta Prx1}$* chondrocytes. (F) Percentage of MBD2-mediated enrichment in methylated DNA in 1 μg total DNA from *Dnmt1$^{flox}$* (n=6) and *Dnmt1$^{\Delta Prx1}$* (n=6) chondrocytes. (G) Peak count frequency of MBD-Seq. (H) Methylated DNA signals for selected genes as illustrated with the Integrative Genome Viewer (IgV). All data are mean ± s.d. **** $p < 0.0001$ (unpaired t test).

### Figure 5 - figure supplement 5

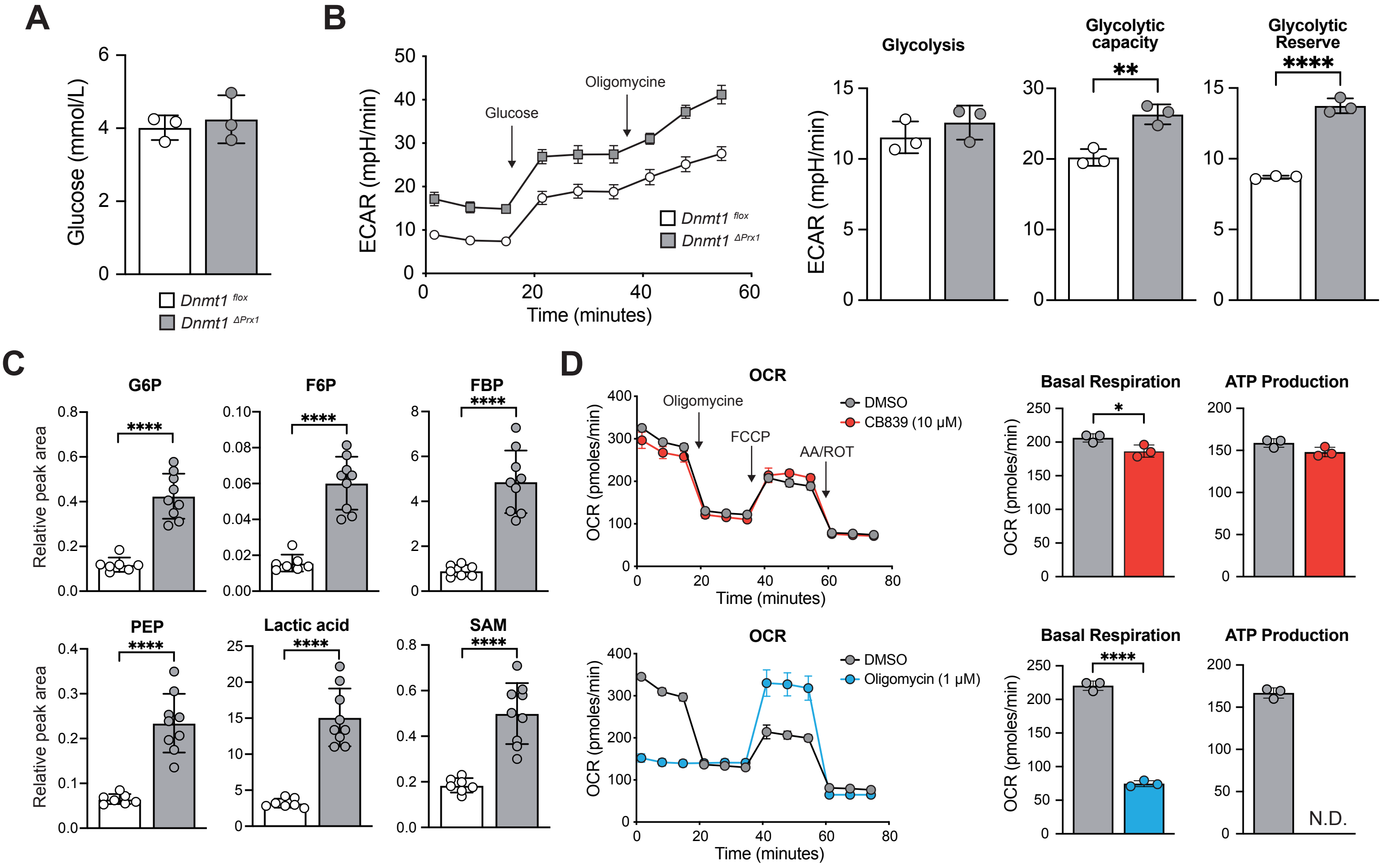


**Figure S5. Increased glycolysis metabolism in Dnmt1-deficient chondrocytes**

(A) Glucose uptake in *Dnmt1*$^{flox}$ (n=3) and *Dnmt1*$^{\Delta Prx1}$ (n=3) chondrocytes. (B) Left panel: Extracellular acidification rate (ECAR) assessed after addition of glucose and oligomycin at the indicated times. Right panel: Glycolysis, glycolysis capacity, and glycolytic reserve measured in *Dnmt1*$^{flox}$ (n=3) and *Dnmt1*$^{\Delta Prx1}$ (n=3) chondrocytes. (C) Quantification of relative levels of energy metabolites by metabolome analysis of *Dnmt1*$^{flox}$ (n=7) and *Dnmt1*$^{\Delta Prx1}$ (n=9) chondrocytes. (D) Confirmation of OCR inhibition in *Dnmt1*$^{\Delta Prx1}$ chondrocytes treated with energy metabolism inhibitors (CB839: Glutaminase inhibitor (n=3), Oligomycin: ATP synthesis inhibitor (n=3)). All data are mean ± s.d. * $p < 0.05$, ** $p < 0.01$, **** $p < 0.0001$ (unpaired *t* test).

### Figure 6 - figure supplement 6

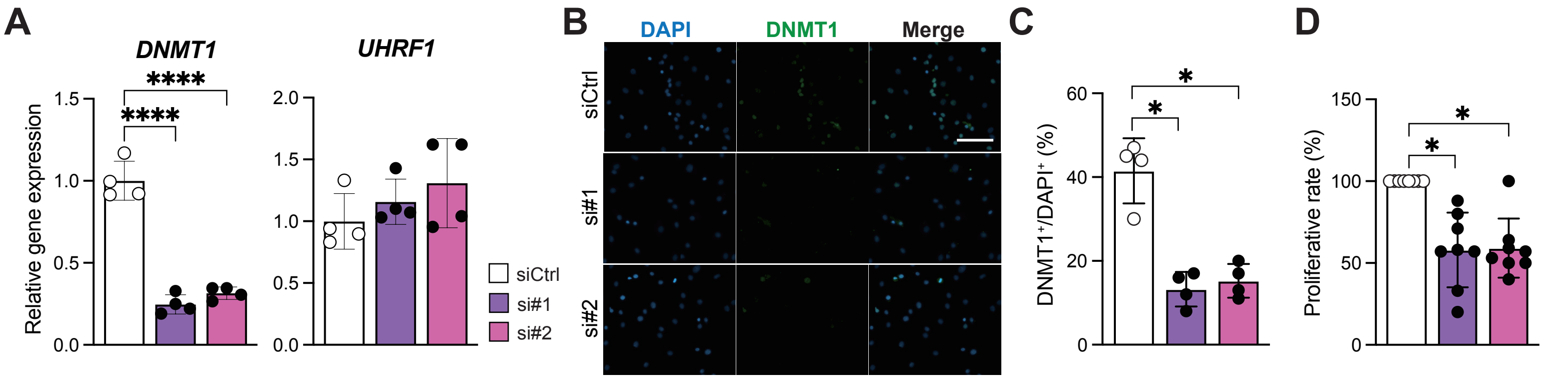


**Figure S6.**
(A) Expression levels of genes related to maintenance of DNA methylation after differentiation stimulus by BMP2 as determined by real-time RT-PCR (n=4). (B) Immunocytochemistry for DNMT1 (Green) and DAPI (Blue) in human articular chondrocytes. (C) Quantification of DNMT1 expression in human articular chondrocytes (n=4). (D) Cell proliferation rate calculated by cell counting (n=8). All data are mean ± s.d. * $p < 0.05$, ** $p < 0.01$ (Tukey's post-hoc test).

**Supplementary Table 1.**
**Summary of the primers and oligonucleotide sequences in this study.**

Primers for qPCR (Mouse)

| | Forward | Reverse |
|---|---|---|
| mAlp | ACACCTTGACTGTGGTTACTGCTGA | CCTTGTAGCCAGGCCCGTTA |
| mCol10a1 | GGGACCCCAAGGACCTAAAG | GCCCAACTAGACCTATCTCACCT |
| mCol1a1 | ACATGTTCAGCTTTGTGGACC | TAGGCCATTGTGTATGCAGC |
| mCol2a1 | AGGGCAACAGCAGGTTCACATAC | TGTCCACACCAAATTCCTGTTCA |
| mDnmt1 | AAGAATGGTGTTGTCTACCGAC | CATCCAGGTTGCTCCCCTTG |
| mGapdh | AGGTCGGTGTGAACGGATTTG | TGTAGACCATGTAGTTGAGGTCA |
| mMmp13 | GGTCCTTGGAGTGATCCAGA | TGATGAAACCTGGACAAGCA |
| mOcn | ACAGACTCCGGCGCTACCTT | AATAGTGATACCGTAGATGCGTTTG |
| mOpn | TACGACCATGAGATTGGCAGTGA | TATAGGATCTGGGTGCAGGCTGTAA |
| mRunx2 | GACGTGCCCAGGCGTATTTC | AAGGTGGCTGGGTAGTGCATTC |
| mSox9 | AGCTCACCAGACCCTGAGAA | TCCCAGCAATCGTTACCTTC |
| mSp7 | CTTCCCAATCCTATTTGCCGTTT | CGGCCAGGTTACTAACACCAATCT |
| mTet1 | CAGCCGTTGAAATACATGCTC | ACATCCCACAGACCGAAGA |
| mTet2 | AGAGCCTCAAGCAACCAAAA | ACATCCCTGAGAGCTCTTGC |
| mTet3 | CCTTTTCTCCATACCGATCCTC | GAGTTCCCTACCTGCGATTG |
| mUhrf1 | CCACACCGTGAACTCTCTGTC | GGCGCACATCATAATCGAAGA |

Primers for qPCR (Human)

| | Forward | Reverse |
|---|---|---|
| hACAN | ACTCTGGGTTTTCGTGACTCT | ACACTCAGCGAGTTGTCATGG |
| hALPL | ACCACCACGAGAGTGAACCA | CGTTGTCTGAGTACCAGTCCC |
| hBGLAP | CACTCCTCGCCCTATTGGC | CCCTCCTGCTTGGACACAAAG |
| hCOL10A1 | ATGCTGCCACAAATACCCTTT | GGTAGTGGGCCTTTTATGCCT |
| hCOL2A1 | TGGACGATCAGGCGAAACC | GCTGCGGATGCTCTCAATCT |
| hDNMT1 | CCTAGCCCCAGGATTACAAGG | ACTCATCCGATTTGGCTCTTTC |
| hGAPDH | ACAACTTTGGTATCGTGGAAGG | GCCATCACGCCACAGTTTC |
| hMMP13 | ACTGAGAGGCTCCGAGAAATG | GAACCCCGCATCTTGGCTT |
| hNOCT | TCTCGCCAAGACACTGAACAG | GGCCCTGCATTCCTCAAGAAG |
| hRAB3GAP1 | GCCTGGTAAGATGGTATGGGC | GCACTTAGATTCGCTGAGAACA |
| hRUNX2 | TGGTTACTGTCATGGCGGGTA | TCTCAGATCGTTGAACCTTGCTA |
| hSIRT5 | GCCATAGCCGAGTGTGAGAC | CAACTCCACAAGAGGTACATCG |
| hSLC38A6 | GCTTTTGACAGTCCCTCTAATCC | TTGATGTACTGGCACCAACTAC |
| hSOX9 | AGCGAACGCACATCAAGAC | CTGTAGGCGATCTGTTGGGG |
| hSP7 | CCTCTGCGGGACTCAACAAC | AGCCCATTAGTGCTTGTAAAGG |
| hUHRF1 | AGGTGGTCATGCTCAACTACA | CACGTTGGCGTAGAGTTCCC |
